# Mapping the task-general and task-specific neural correlates of speech production: meta-analysis and fMRI direct comparisons of category fluency and picture naming

**DOI:** 10.1101/2023.09.27.559692

**Authors:** Gina F. Humphreys, Matthew A. Lambon Ralph

**Affiliations:** MRC Cognition & Brain Sciences Unit, University of Cambridge, Cambridge, UK

**Keywords:** Speech production, category fluency, picture naming, fMRI, meta-analysis

## Abstract

Improving our understanding of the neural network engaged by different forms of speech production is a crucial step for both cognitive and clinical neuroscience. We achieved this aim by exploring two of the most commonly utilised speech production paradigms in research and the clinic, which have been rarely, if ever, compared directly: picture naming and category fluency. This goal was achieved in this two study investigation through a full ALE meta-analysis as well as a targeted fMRI study. Harnessing the similarities and differences between the two tasks offers a powerful methodology to delineate the core systems recruited for speech production, as well as revealing task-specific processes. The results showed that both tasks engaged a bilateral fronto-temporal speech production network, including executive and motor frontal areas, as well as semantic representational regions in the ATL, bilaterally. In addition, it was found that the extent of relative frontal lateralisation was task-dependent with the more executively-demanding category fluency task showing augmented left hemisphere activation. The results have implications for neurocomputational speech production models and the clinical assessment of speech production impairments.

Open access: For the purpose of open access, the UKRI-funded authors have applied a Creative Commons Attribution (CC BY) licence to any Author Accepted Manuscript version arising from this submission.

## Introduction

Speech production is phenomenal human capacity. Yet, despite decades of neuropsychological and imaging research it is still unclear how the brain transforms a non-verbal thought into speech output. Here we directly investigate this issue using two of the most commonly utilised speech production paradigms in research and the clinic: picture naming and category fluency. Indeed, harnessing the similarities and differences between tasks offers a powerful methodology to delineate the core language and cognitive systems recruited for speech production, as well as revealing task-specific processes. In terms of neurocognitive processing, picture naming and category fluency share core similarities that are central to speech production (e.g., the areas recruited for semantics, phonology and motor output) but nevertheless there are important task differences. Picture naming is driven by bottom-up visual input, therefore in addition to core-production regions it relies largely on the visual object recognition system. In contrast, verbal fluency requires little/no visual processing but is instead driven by multiple top-down demands on the executive systems, including linguistic and non-linguistic regions involved working memory, search mechanisms, and the inhibition of inappropriate or already produced items.

This work has novelty and importance for three major reasons. First, from a basic science point of view, picture naming and category fluency are commonly used as proxies to examine the neural systems engaged by production, and have had a major influence neurocognitive models of speech production. Secondly, from a clinical perspective, one or both tasks are frequently used as a proxy for speech production as part of a generalised assessment in a neurological clinic, as well as to guide resection during neurosurgery. Despite their prevalent clinical use, we currently do not have a good understanding of their neural basis. Thirdly, whilst category fluency and picture naming are frequently used tasks in fMRI, from a methodological perspective, existing neuroimaging paradigms are frequently suboptimal at detecting whole-brain effects, therefore potentially missing key production regions. Achieving full-brain coverage is important not only for developing fully specified neuroanatomical models but especially if one is to use fMRI as a neurosurgical tool for pre-operative planning. Here we conducted the first direct comparison of picture naming and category fluency tasks via two major investigations: 1) a meta-analysis of past neuroimaging studies; and 2) an fMRI study to directly compare activation patterns across tasks using a signal-optimised fMRI paradigm.

### Unresolved issues within neurocognitive theory

Neurocognitive theories tend to emphasise the importance of a left-lateralised dorsal network for speech production, involving posterior temporo-parietal cortex and inferior frontal gyrus (IFG) connected via the arcuate fasciculus. This was first highlighted in classical models, for instance, Geschwind (1965), influenced by earlier work by Broca and Wernicke, proposed that semantic content stored in the angular gyrus is sent via the arcuate to the IFG, where meaning is converted to speech output (though recent re-visits of the classical literature (Weiller, Bormann, Saur, Musso, & Rijntjes, 2011) suggest that Wernicke actually considered the ventral pathway as dominant for production). A similar dorsal route is highlighted in many contemporary neurocognitive models; a range of theories suggest that lexical-semantic and phonological access is achieved via the posterior temporal-parietal cortex, following which information is transferred to the IFG via the arcuate fasciculus whereby phonological information is transformed into articulatory code for motor output (Friederici, 2002; Hagoort, 2005; Hickok, 2012; Hickok, Houde, & Rong, 2011; Hickok & Poeppel, 2000, 2004, 2007; Indefrey & Levelt, 2004). According to these models, this so called “dorsal” route is in contrast to a temporal lobe “ventral” route which and is critical for receptive speech (Friederici, 2002; Hagoort, 2005; Hickok, 2012; Hickok et al., 2011; Hickok & Poeppel, 2000, 2004, 2007; Indefrey & Levelt, 2004).

### The anterior temporal lobe (ATL)

In some versions of the dual-stream model, ventral areas such as the ATL, have no/minimal involvement in production (Hickok, 2012; Hickok et al., 2011; Hickok & Poeppel, 2000, 2004, 2007). These versions of the dual-stream model are heavily influenced by findings from the neuroimaging literature. For instance, a large review showed reliable fronto-posterior temporal activation for speech production tasks using PET/fMRI or EEG/MEG (Indefrey, 2011; Indefrey & Levelt, 2004). The patient literature, however, paints a different picture. Here, damage to the ventral pathway, specifically to the ATL in semantic dementia (SD) or temporal lobe resection, results in a pronounced deficit in meaningful speech production and comprehension, as well as non-verbal semantic tasks (Lambon Ralph, Ehsan, Baker, & Rogers, 2012; Lambon Ralph, Jefferies, Patterson, & Rogers, 2017). This is in keeping with the proposal that the ATLs act as a semantic hub that supports transmodal conceptual representation, and is critical to speech production (Jefferies & Lambon Ralph, 2006; Lambon Ralph et al., 2012; Lambon Ralph et al., 2017; Lambon Ralph, McClelland, Patterson, Galton, & Hodges, 2001). Indeed, this theory has gained converging support across a variety of methodologies, including TMS, fMRI, ECoG, as well as from computational modeling (Lambon Ralph et al., 2017; Lambon Ralph, Sage, Jones, & Mayberry, 2010; Pobric, Jefferies, & Lambon Ralph, 2010; Rogers et al., 2021; Sato et al., 2021; Shimotake et al., 2014; Woollams, Lindley, Pobric, & Hoffman, 2017). Alternative dual-route language frameworks that incorporate an ATL-semantic pathway, implemented in computational models (Ueno, Saito, Rogers, & Lambon Ralph, 2011), demonstrate key divisions of labour across the dorsal and ventral pathways with the latter crucial for semantically-driven speech production (unlike repetition in the dorsal pathway). If true, one should expect that the ATL be engaged by both picture naming and category fluency tasks.

### The angular gyrus (AG)

It has been proposed that the AG acts as a multi-modal semantic storage hub, similar in function to the ATL (Binder, Desai, Graves, & Conant, 2009; Geschwind, 1965), based largely on neuroimaging evidence showing that the AG is one of the most reliably engaged regions by contrasts of word>non-words or concrete>abstract (Binder et al., 2009). Nevertheless, the majority of evidence comes from receptive rather than expressive language tasks. Those tasks that have included speech production, have not consistently found AG engagement and recent work suggests that the minority of positive production tasks might relate to autobiographical recall (Geranmayeh et al., 2012; Humphreys, Halai, Branzi, & Lambon Ralph, 2022).

### The lateral front cortex

In the standard dual-stream framework, the lateral frontal cortex is primarily considered to be involved in articulatory motor planning (Hickok, 2012; Hickok et al., 2011). Yet the lateral frontal cortex is a heterogeneous region packed with multiple additional functions, such a phonological and executive processes. In terms of executive mechanisms, dorsal lateral prefrontal cortex is involved in domain-general executive control and forms a key part of the multiple demand (MD) network (Fedorenko, Duncan, & Kanwisher, 2013), and the IFG forms part of the semantic control network, a network involved in the use and manipulation of semantic information (Jefferies & Lambon Ralph, 2006; Lambon Ralph et al., 2017). In terms of language processing, damage to the IFG can impair performance in executively demanding expressive and receptive semantic tasks, (Jefferies & Lambon Ralph, 2006; Noonan, Jefferies, Corbett, & Lambon Ralph, 2010; Rogers, Patterson, Jefferies, & Lambon Ralph, 2015), and similar results have been found using TMS in healthy participants (Krieger-Redwood & Jefferies, 2014; Whitney, Kirk, O’Sullivan, Lambon Ralph, & Jefferies, 2010). Likewise, recent work with continuous speech production has shown a key role for ventral IFG in semantically coherent speech production (Hoffman, 2019; Hoffman, Loginova, & Russell, 2018). We therefore predicted that the left lateral frontal cortex plays an important role in speech production beyond articulatory motor planning.

### Unresolved issues with clinical implications

#### Language laterality

Clinically, language production is frequently considered a largely left hemisphere function (Eggert, 1977; Geschwind, 1972; Lichtheim, 1885). Nevertheless, with advancements in functional imaging there is increasing evidence of a bilateral albeit asymmetric neuroimaging pattern of activation in speech production (Blank, Scott, Murphy, Warburton, & Wise, 2002; Geranmayeh et al., 2012; Humphreys, Halai, et al., 2022; Moore & Price, 1999; Zelkowicz, Herbster, Nebes, Mintun, & Becker, 1998). To further muddy the waters, the extent of left vs. bilateral activation appears task-dependent (Birn et al., 2010; Blank et al., 2002; Geranmayeh et al., 2012; Moore & Price, 1999; Vitali et al., 2005; Zelkowicz et al., 1998). The extent to which speech production lateralises has implications for patient treatment, for instance, the decision to assess language function during neurosurgery and whether or not to refer a patient to speech and language therapy.

#### Speech assessment in patients

Picture naming and/or category fluency are often used as part of a generalised neuropsychological assessment in the clinic, as well as to guide resection during neurosurgery. These tasks are very practical in the clinic since they are quick to perform and require relatively little technology, and thus make an attractive task for language screening. Issues may arise, however, if only one task is used in isolation as a generalised proxy for speech production, as patients can show varying performance across measures, for instance patients may be more severally impaired in category fluency assessments compared to picture naming due its enhanced executive demands (Noonan et al., 2010; Robinson, Blair, & Cipolotti, 1998; Schumacher, Halai, & Lambon Ralph, 2022). Indeed, given its multiple cognitive demands, it is entirely possible to have poor category and letter fluency scores as a result of entirely non-language impairments (Henderson, Peterson, Patterson, Lambon Ralph, & Rowe, 2022; Hodges & Patterson, 1995).

### Methodological issues

There are a number of key methodological limitations in the current speech production literature. First, as mentioned above, the ATL is highly susceptible to signal dropout using conventional fMRI paradigms (Halai, Welbourne, Embleton, & Parkes, 2014) and is much more likely to be observed when contrasted against an active task baseline than rest (Humphreys, Hoffman, Visser, Binney, & Ralph, 2015; Visser, Jefferies, & Lambon Ralph, 2010). These and other factors mean that the current fMRI speech production literature may well be missing the ATL from the full production network. Indeed, ATL activation has been observed during picture naming tasks using PET (Blank et al., 2002; Moore & Price, 1999; Zelkowicz et al., 1998), a technique that does not suffer from signal dropout. Achieving full-brain coverage is important not only for developing neuroanatomical models but especially if one is to use fMRI with patients to assess the functional integrity of the ventral language pathways (e.g., in aphasia or as a neurosurgical tool for pre-operative planning: (Rice, Caswell, Moore, Hoffman, & Lambon Ralph, 2018; Robson et al., 2014)). Secondly, existing imaging paradigms often use covert rather than overt speech production which will limit the extent of the observed the production network. Thirdly, existing studies tend to suffer from a lack of experimental control. For instance, category fluency studies often use very long blocks and do not take into account the speech rate or the number of words produced.

### Aims

Here we conducted an investigation containing two linked, full studies: (1) a meta-analysis of the existing picture naming and category fluency literature, as well as (2) the first within-subject fMRI study to compare the tasks directly. Meta-analyses are powerful techniques to identify reliable findings from the existing literature that hold across investigations and are independent of study-specific factors and other idiosyncrasies. By their very nature of relying on the extant literature, however, meta-analytic results can be biased by any methodological and sampling issues that systematically occur across studies (of which there are several in this case). Hence, combining the advantages of meta-analyses with those from a well-controlled within-subject fMRI study is the optimal approach. In our fMRI study we controlled for these methodological issues by: 1) using a dual-echo fMRI protocol which improves ATL signal detection (Halai et al., 2014); 2) using overt speech production tasks; and 3) match the rate and the number of words produced across tasks by using a paced paradigm.

We had three key aims: 1) the use of two very different speech production methodologies allow greater precision when determining the shared cognitive mechanism and associated brain regions across tasks using conjunction analyses (e.g., semantic, phonological, and motor components), and also to delineate task-specific processes through direct task contrasts; 2) we assessed the extent to which the production network is left lateralised (vs. bilateral), and whether the extent of lateralisation is task-dependent; and 3) we determined the role of the ATL, lateral frontal cortex and AG in speech production using targeted ROI analyses.

## Methods

### Meta-analysis

#### Study selection

The picture naming and category-fluency studies included in the meta-analysis were identified by keyword searches Google Scholar (these included “Speech production” or “Word production” or “Naming’ or “Fluency” AND “fMRI” or “PET”) resulting in the identification of over 500 studies. Of those studies, only those including either/both a picture naming and category-fluency task and reported peak activation in standard space (Talairach or MNI) based on whole-brain statistical comparisons were included. This resulted in the identification of 62 picture-naming studies (628 foci), and 44 category-fluency studies (407 foci).

#### ALE analyses

The ALE analyses were carried out using GingerAle 3.0.2 (Eickhoff et al., 2009; Laird et al., 2005). All activation peaks were converted to MNI standard space using the built-in Talairach to MNI (SPM) GingerALE toolbox. Analyses were performed with voxel-level thresholding at a p-value of 0.001 and cluster-level FWE-correction with a p-value of 0.05 over 10,000 permutations.

### fMRI study

#### Participants

Twenty participants took part in the fMRI study (average age = 24.30, SD = 3.96; N female = 12). All participants were native English speakers with no history of neurological or psychiatric disorders and normal or corrected-to-normal vision.

#### Task design and procedures

There were three experimental tasks presented using a randomised blocked-design. At the beginning of the block the participants were cued with a task instruction for a duration of 1.5 seconds. Four items were presented per block, each for two seconds separated by a 25ms fixation cross (block length = 10.5 seconds). There were 80 items in each task (20 blocks per condition). The blocks were randomly separated with 30 rest blocks consisting of a fixation cross.

#### Picture naming task

The participants were cued to “Name the picture” at the beginning of each block. The pictures consisted of black-and-white line drawing from the Snodgrass (Snodgrass & Vanderwart, 1980) picture set and consisted of a mixture of living and non-living items. The participants were instructed to name the picture before it left the screen.

#### Category fluency task

At the beginning of each block the participants were cued with a semantic category (e.g. “Name zoo animals”). During each trial a scrambled image was presented on the screen. The participants were instructed to generate a word every time a scrambled image was presented. This provided a method of pacing the rate of speech production to match the picture naming task. The scrambled picture also acted as a low-level visual control for the pictures in the picture naming task.

#### Control task

On each trial the participants were cued to “Say OK” overtly when a scrambled line drawing was centrally presented. The control trials acted as a control for motor aspects of speech production as well as low level visual processing in the experimental tasks.

#### Task acquisition parameters

Images were acquired using a 3T Philips Achieva scanner using a dual gradient-echo sequence, which has improved signal relative to conventional techniques, especially in areas associated with signal loss (Halai AD et al. 2014). 31 axial slices were collected using a TR = 2.8 seconds, TE = 12 and 35ms, flip angle = 95°, 80 x 79 matrix, with resolution 3 x 3 mm, slice thickness 4mm. Across all tasks, 359 volumes were acquired in total collected in one run of 1005.2 seconds. B0 images were also acquired to correct for image distortion.

### Task data analysis

#### Preprocessing

The dual-echo images were first B0 corrected and then averaged. Data were analysed using SPM12. Images were motion-corrected and co-registered to the participants T1 structural image, and then spatially normalised into MNI space using DARTEL (Ashburner, 2007). The functional images were then resampled to a 3 × 3 × 3mm voxel size and smoothed with an 8mm FWHM Gaussian kernel.

#### General Linear Modelling

The data were filtered using a high-pass filter with a cut-off of 190s and then analysed using a general linear model. At the individual subject level, each condition for each task was modelled with a separate regressor and events were convolved with the canonical hemodynamic response function. Time and dispersion derivatives were added and motion parameters were entered into the model as covariates of no interest (no participants exceeded the absolute motion threshold set at the maximum of one voxel of displacement (Johnstone et al., 2006)). At the individual level, the naming and fluency trials were separately contrasted against the control condition as well as directly against one another. We also contrasted both language tasks > control. A standard voxel height threshold p < .001, cluster corrected using FWE p < .05 was used.

#### ROI analyses

We also conducted targeted ROI analyses. The purpose of these was twofold. Firstly, we compared activation from key left-hemisphere ROIs with their right-hemisphere homologue in order to test for laterality effects across hemispheres. Secondly, we compared task activation within each ROI to test our predictions regarding the functional engagement of these regions. Specifically, we included four lateral frontal ROIs (two left hemisphere and two right hemisphere) in order to assess the extent to which each task engaged areas associated with executive function. Specifically, we included one ROI in the left IFG region associated with semantic control based on the coordinates from meta-analysis of studies with high-vs. low-semantic control demands (Noonan, Jefferies, Visser, & Lambon Ralph, 2013), and one in the left inferior frontal sulcus using the peak coordinate from an atlas of the MD network (https://evlab.mit.edu/funcloc/), as well as their right hemisphere homologues. In order to assess the functional involvement of the ATL across tasks we also included bilateral ventral ATL ROIs based on the coordinates from a semantic study (Binney, Embleton, Jefferies, Parker, & Lambon Ralph, 2010). Finally, we included an ROI of the left AG based on the peak coordinates from meta-analyses of semantic studies (Humphreys & Lambon Ralph, 2015) to test the hypothesis that the left AG acts as an additional semantic hub (Binder et al., 2009).

## Results

### Meta-analysis results

The analysis of category fluency and picture naming studies showed that both tasks reliably engaged overlapping frontal areas (inferior frontal gyrus (IFG), middle frontal gyrus (MFG), premotor and motor cortex, and the supplementary motor area (SMA)/anterior cingulate cortex (ACC)), in areas associated with executive and motor processing (Fedorenko et al., 2013; Hickok, 2012; Lambon Ralph et al., 2017). Significant right hemisphere frontal recruitment was only found for the fluency studies but not picture naming, and these clusters were comparatively smaller in comparison to those in the left hemisphere. Beyond the frontal cortex, category fluency showed very little recruitment, whereas picture naming revealed significant clusters in bilateral visual and posterior ventral temporal cortex (fusiform gyrus) in areas associated with visual object recognition (DiCarlo & Cox; Riesenhuber & Poggio, 1999). (Figure 1, Table S1).

**Figure 1.**
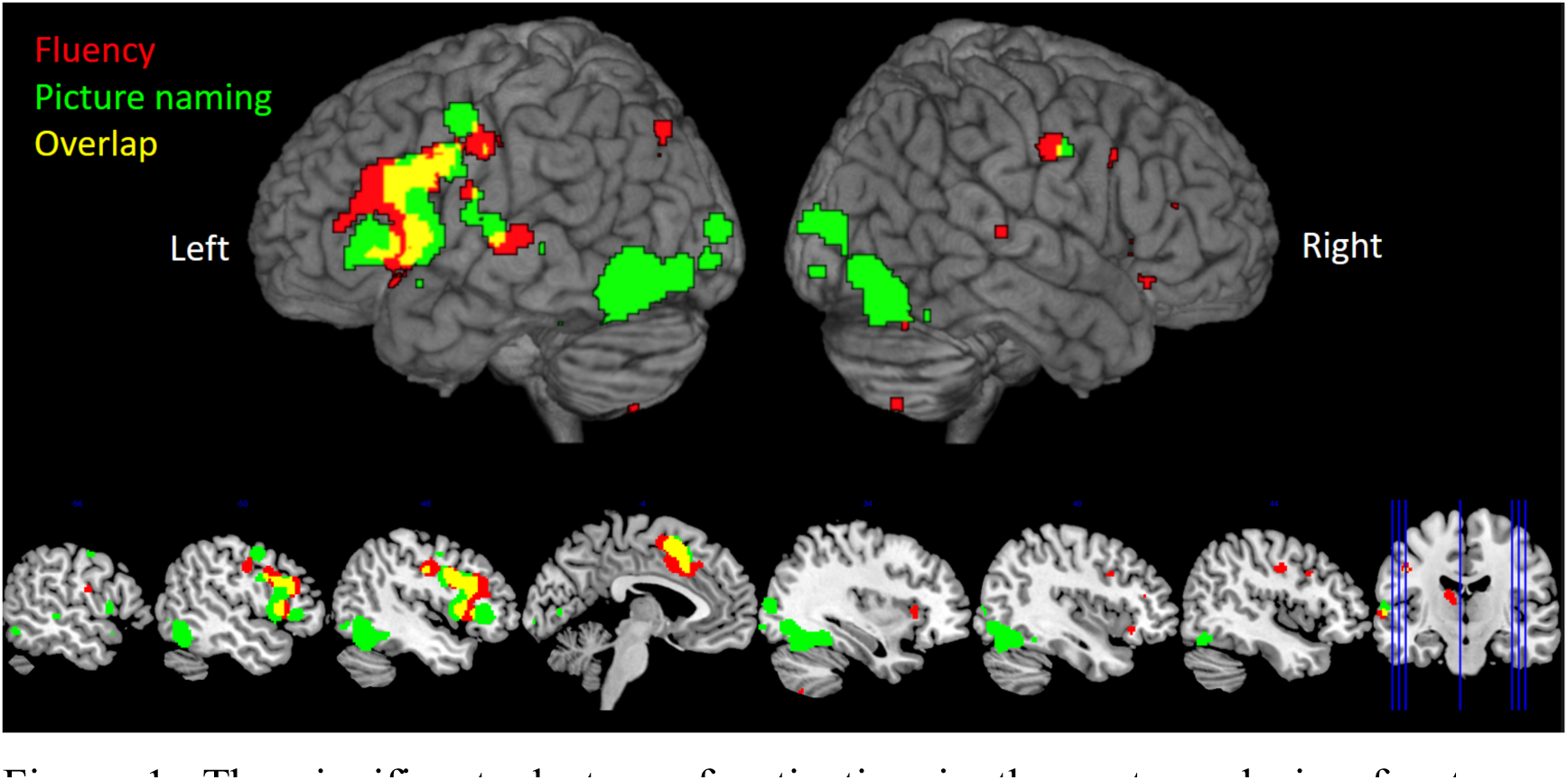
The significant clusters of activation in the meta-analysis of category fluency studies (red) and picture naming (green) (thresholded at a p < 0.001, cluster-level FWE-correction p < 0.05). The overlap is shown in yellow.

A direct comparison of fluency and picture naming results showed significantly stronger recruitment of left frontal (premotor, SMA/ACC), and right frontal cortices (MFG) for the fluency compared to naming tasks, although the left hemisphere clusters were larger. The direct contrast of picture naming and fluency revealed clusters in bilateral visual and posterior ventral temporal cortex (fusiform gyrus) (see Table S1). Neither task was found to reliably activate the ATL. We propose that this may be due to there being limited fMRI signal in this area (Visser et al., 2010) - indeed studies that used PET rather than fMRI did find ATL involvement (Blank et al., 2002; Moore & Price, 1999; Zelkowicz et al., 1998). Indeed, the results of the fMRI study that used a technique for improved ATL signal detection support this conclusion (see next).

### fMRI results

#### The core production network

To determine a task-general network we examined the areas of overlap between the Fluency > control and Naming > control contrasts (Figure 2, Table S2). A common pattern of activation was observed in bilateral frontal areas that are associated with executive and motor processes (inferior frontal gyrus, middle frontal gyrus, the supplementary motor area, anterior cingulate cortex, and pre-motor and primary-motor cortices) (Binder et al., 2009; Fedorenko et al., 2013; Hickok, 2012; Jackson, 2021). Of note, bilateral ventral ATL (anterior fusiform gyrus) was also engaged by both tasks indicating common involvement of the semantic hub across tasks (Lambon Ralph et al., 2017). In addition to this fronto-temporal network, common activation was observed in bilateral superior parietal areas previously associated with domain-general executive control and visuo-spatial processes (Fedorenko et al., 2013; Kravitz, Saleem, Baker, & Mishkin, 2011), as well occipital cortex.

**Figure 2.**
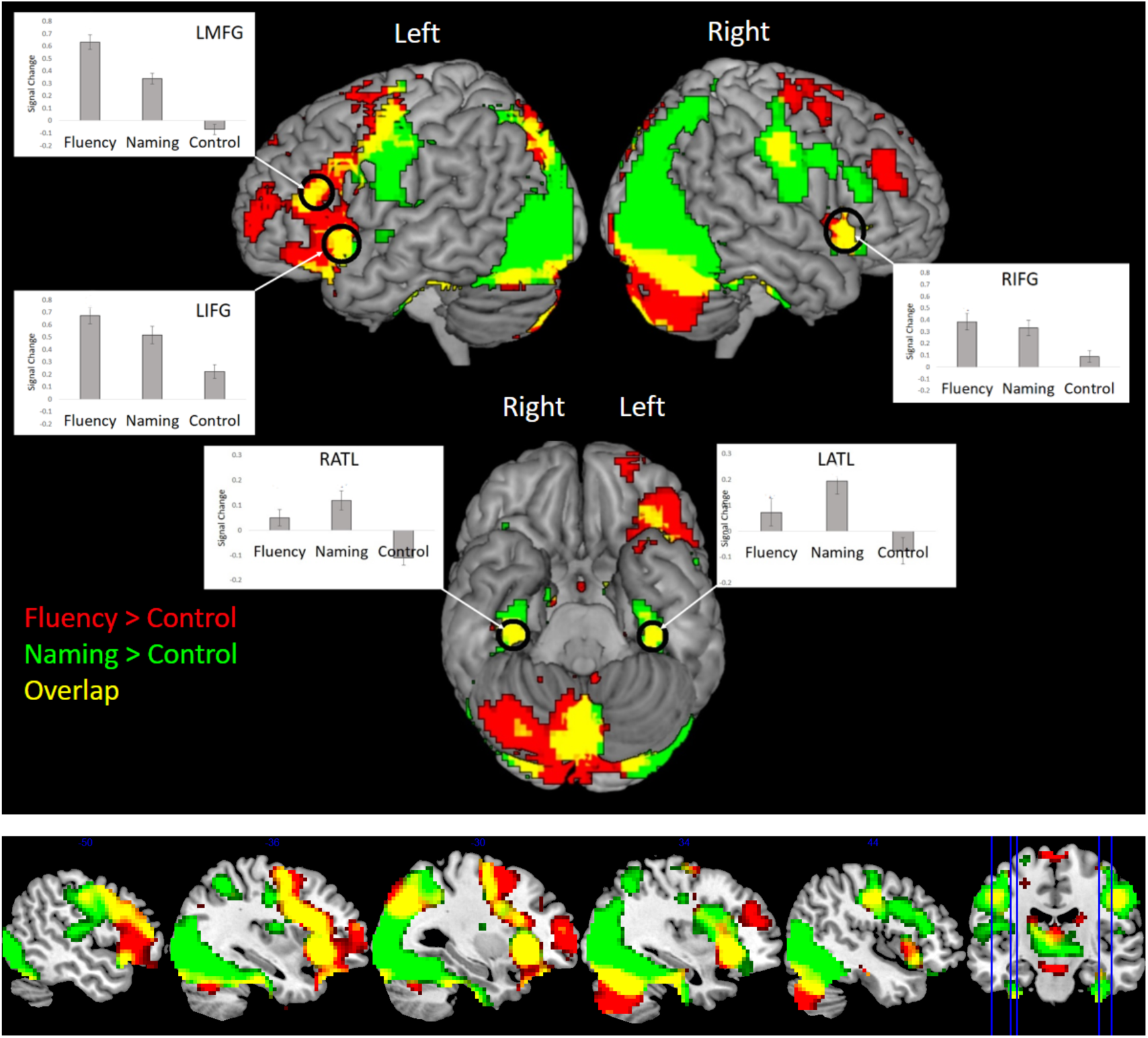
The results from the fMRI study for the contrasts Fluency > Control (red) and Naming > Control (green) (thresholded at p < .001, cluster corrected using FWE p < .05). The overlap between each network is shown in yellow.

#### Task-varying networks

Fluency > Naming: Whilst large areas of the left frontal cortex was shown to be engaged by both fluency and picture naming, activation was found to be stronger in many areas for the fluency task compared to naming (IFG, MFG, SMA, ACC, and premotor cortex). This enhanced frontal activation is consistent with the increased executive demands of the fluency compared to naming task. This was also true in right dorso-medial frontal cortex (ACC and MFG), although these clusters were comparatively smaller compared to the left hemisphere and were restricted to dorso-lateral regions associated with domain-general executive control. Whilst IFG is frequently associated with semantic control (Jackson, 2021), the area immediately dorsal shows a more domain-general function (Chiou, Jefferies, Duncan, Humphreys, & Lambon Ralph, 2023). Outside the frontal lobe, the only region to show a fluency > naming difference was the left AG. Since the AG was not revealed in the contrast of Fluency > control (even at a very lenient threshold of p < .05, uncorrected) it may not play a central role in Fluency per se (indeed, see ROI results below for further investigation of the left AG). All results are shown in Figure 3 and Table S2.

**Figure 3.**
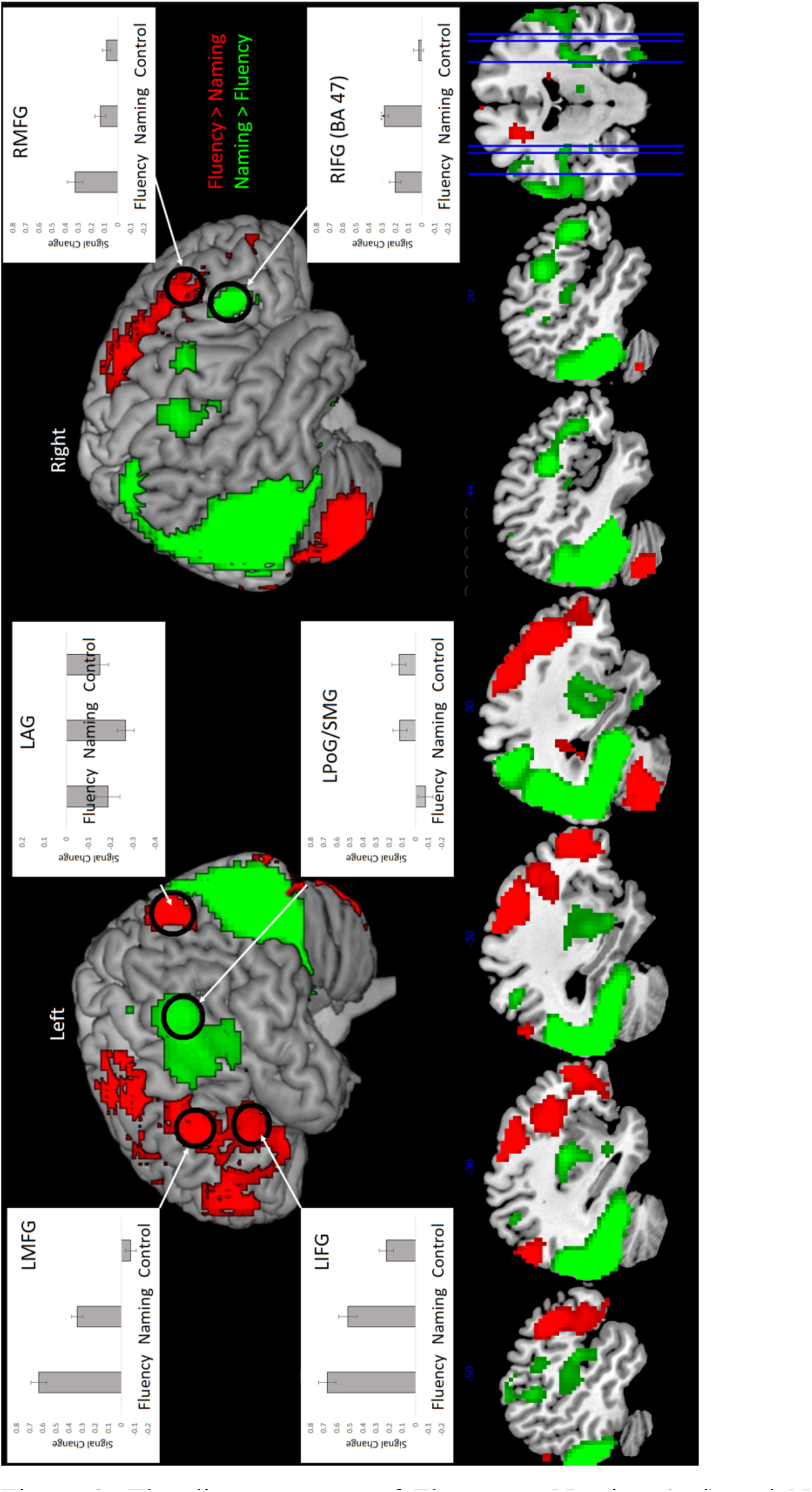
The direct contrast of Fluency > Naming (red) and Naming > Fluency (green) (thresholded at p < .001, cluster corrected using FWE p < .05).

Naming > Fluency: As expected, picture naming elicited stronger activation in the bilateral occipital cortex as well occipito-temporal activation along the length of fusiform gyrus, in areas associated with visual-object recognition (DiCarlo & Cox; Riesenhuber & Poggio, 1999). Additionally, in the frontal lobe, whilst the left frontal cortex was more strongly engaged by fluency, picture naming showed stronger activation for the right IFG and premotor cortex (refer to the ROI results for further investigations into the laterality differences). Unexpectedly, naming was also found to show stronger activation of bilateral postcentral/supramarginal gyrus. This area was not identified by the contrast of naming > control, even at lenient threshold of p < .05 uncorrected. Therefore, whilst the explanation of this result is unclear, this region does not appear to play an important role in picture naming per se. All results are shown in Figure 3 and Table S2.

#### ROI analyses

Whilst language production is classically considered exclusive to the left hemisphere, the imaging literature suggests that there may be a more bilateral pattern, as well as the possibility that the extent of laterality might be task-dependent (Blank et al., 2002; Stefaniak, Geranmayeh, & Lambon Ralph, 2022; Stefaniak, Halai, & Lambon Ralph, 2020). Planned ROI analyses were conducted to examine variations in laterality across tasks in three key regions of the executive and network: including 1) the semantic hub region of the vATL, as well as two lateral frontal executive areas: 2) the semantic control region within IFG (Jackson, 2021) and, 3) the more dorsal MD control region of IFS (Fedorenko et al., 2013) (Figure 4).

**Figure 4.**
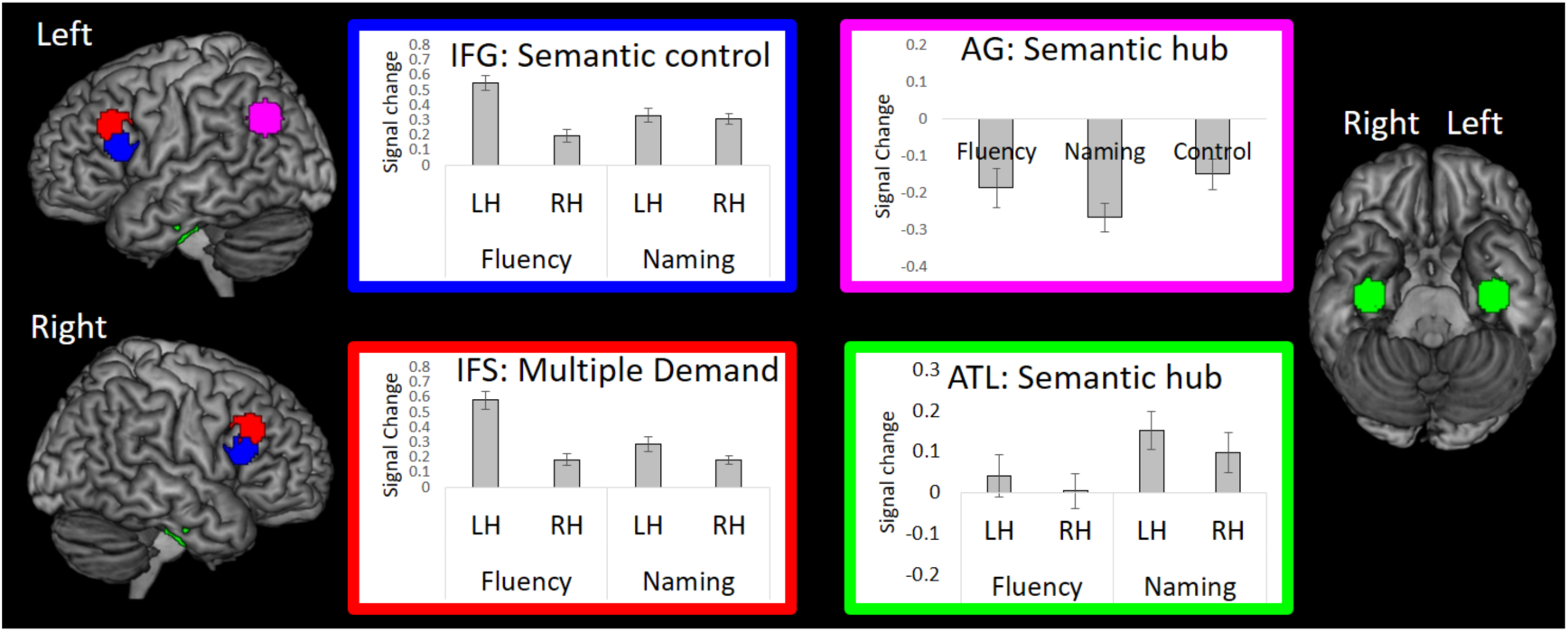
The results from the ROI analysis.

Repeated measures ANOVAs were used to test for laterality x task effects in each ROI. The semantic control region of the IFG was found to show a significant effect of hemisphere (F(19) = 26.72, p < .001), no effect of task (F(19) = 2.56, p = .13), and a significant hemisphere x task interaction (F(19) = 112.05, p < .001). Pairwise comparisons showed that the interaction was explained by significantly increased left hemisphere compared to right hemisphere activation for the fluency task (t(19) = 9.41, p < .001), but no hemispheric difference for the naming task (t(19) = 0.59, p = .28). This suggests that left hemisphere activation is boosted for the fluency task whereas the naming task shows overall weaker but bilateral activation. This result is consistent with outcome of the whole-brain analysis.

The MD region of the IFS showed a similar pattern as IFG: a significant effect of hemisphere (F(19) = 39.56, p < .001), and a significant hemisphere x task interaction (F(19) = 62.22, p < .001). But additionally showed a significant effect of task (F(19) = 15.86, p < .001). Pairwise comparisons showed that the domain-general control region was more engaged by fluency compared to naming in the left hemisphere (t(19) = 6.64, p < .001) and also the right hemisphere (t(19) = 2.55, p < .02). This is consistent with the whole-brain results where bilateral dorsal frontal areas were engaged more strongly by fluency than naming. In addition, whilst the left hemisphere was generally more strongly engaged than the right hemisphere by both the fluency task (t(19) = 7.59, p < .001) and the naming task (t(19) = 3.12, p < .005), the hemispheric difference was significantly greater for the fluency task compared to the naming task (t(19) = 7.89, p < .001). Again, this suggests that left hemisphere activation is particularly boosted for the fluency task, which carries greater executive demand.

For the vATL, there was a moderately significant effect of task F(19) = 8.50, p < .01), but no effect of hemisphere (F(19) = 1.38, p = .25), and no significant hemisphere x task interaction (F(19) = 0.21, p < .70). This indicated whilst there was significantly greater vATL activation overall for the naming vs. fluency task (t(19) = 2.92, p < .01), the pattern of activation was bilateral for both tasks, and is consistent with notions of the vATL operating as a bilateral semantic system (Jung & Lambon Ralph, 2016; Rice, Lambon Ralph, & Hoffman, 2015).

In addition to the hemisphere x task interaction ROI analyses, we also examined the role that the left AG played in each task relative to control (Figure 4). It has been proposed that the AG acts an additional semantic hub (Binder et al., 2009) and hence should be recruited by category fluency and picture-naming. A one-way ANOVA was conducted including naming, fluency, and the control as variables and no significant effect of task was found F(19) = 2.38, p < .11). Furthermore, unlike IFG, IFS and ATL which exhibited positive engagement in the two production tasks (all ts > 4.75, ps < .001, except the left ATL and right ATL for the fluency task (ts < 1.36, P >.05), the AG showed a task-general deactivation (all ts > -3.57, p < .001).

## Discussion

In this double-study investigation of speech production, we combined meta-analysis and fMRI to examine two of the most commonly utilised speech production paradigms in research and the clinic: picture naming and category fluency. We had three key aims. First, to use two very different speech production paradigms to determine what regions are commonly engaged across tasks, and also delineate task-specific components by directly contrasting the tasks. The results showed that both tasks engaged a shared fronto-temporal speech production network, including executive and motor frontal areas, as well as semantic representational regions in the ATL. This network presumably reflects the core speech production systems. Task-specific differences were also revealed, with greater engagement of the frontal executive system for the fluency task, and greater ventral occipito-temporal activation for naming. The second aim of the current study was to test the assumption that the production network is largely left lateralised (vs. bilateral), and the extent to which this is task-dependent. The results showed that both tasks engaged a bilateral fronto-temporal network, however, in lateral frontal areas fluency was associated with boosted left hemisphere activation (Figure 3 and Figure 4), whereas naming showed an overall weaker but more bilateral pattern (Figure 4). The third aim was to investigate the contribution of targeted ROIs to speech production tasks, including frontal areas involved in semantic control and domain-general executive control (IFG, and IFS), as well as proposed semantic hub areas (ATL and AG). Both tasks were found to engage lateral frontal executive regions, although activation was stronger for the fluency compared to naming task, especially in the left hemisphere. The ATL showed bilateral engagement for fluency and naming tasks (although stronger activation for naming), in contrast no evidence was found for the involvement of the AG in either task (in fact, both tasks showed AG deactivation that was equal to or greater than the control task) (Figure 4).

### The core speech production network

Across the two very different speech production tasks we identified a core fronto-temporal production system. This comprised of bilateral frontal cortex in regions associated with motor function and executive control (Fedorenko et al., 2013; Hickok, 2012; Lambon Ralph et al., 2017), and the bilateral ATL associated with semantic representation (Lambon Ralph et al., 2017). How do these findings compare to prominent speech production models, such as the dual-stream model (Hickok, 2012; Hickok et al., 2011; Hickok & Poeppel, 2000, 2004, 2007) and results from the patient literature?

### A bilateral speech production system

We found evidence of a bilateral yet left biased speech production system (although the degree of lateralisation is to some extent task-dependent – see below). This goes against the traditional assumption that speech production relies only on a left-lateralised system which originates from the neuropsychological literature, whereby chronic aphasia is classically associated with left but not right-hemisphere damage (Eggert, 1977; Geschwind, 1972; Lichtheim, 1885). Indeed, a bilateral speech production network has been shown elsewhere in the neuroimaging literature (Blank et al., 2002; Geranmayeh et al., 2012; Humphreys, Halai, et al., 2022; Moore & Price, 1999; Zelkowicz et al., 1998), and TMS studies using healthy participants have also shown that the right-hemisphere plays an important role in language function (Hartwigsen et al., 2013; Jung & Lambon Ralph, 2016; Lambon Ralph, Pobric, & Jefferies, 2009; Pobric et al., 2010; Woollams et al., 2017). How does one reconcile these apparently contradictory findings with the patient literature? One possibility is that language production is a bilateral but asymmetric system, whereby the left-hemisphere has greater computational capacity compared to the right. Indeed, based on this principle, using computational modelling it has been shown that when the right-hemisphere is damaged there is greater capacity within the left-hemisphere to pick up the additional processing requirements, resulting in only a mild/transient language deficit. In contrast, when the left-hemisphere is damaged there are limited right-hemisphere resources available to support recovery resulting severe aphasia (Chang & Lambon Ralph, 2020). Consistent with this model, patients with right hemisphere damage have been found to show transient language deficits in the acute phase with milder deficits remaining chronically, implying some right-hemisphere contribution in the undamaged system (Gajardo-Vidal et al., 2018). Going beyond classical speech areas, the bilateral activation pattern for the ATLs is consistent with previous patient work, rTMS explorations and formal computational models, implying that the ATL is even more inherently bilateral in nature, with substantial semantic deficits only after bilateral rather than unilateral lesions (Lambon Ralph, Cipolotti, Manes, & Patterson, 2010; Lambon Ralph et al., 2012; Pobric et al., 2010; Rice et al., 2018; Schapiro, McClelland, Welbourne, Rogers, & Lambon Ralph, 2013).

### The lateral frontal cortex

is a packed region that serves multiple functions – motor planning, phonological processing and executive control (Fedorenko et al., 2013; Hickok, 2012; Indefrey & Levelt, 2004; Lambon Ralph et al., 2017). Whilst major neurocognitive models highlight the importance of left frontal areas in articulatory motor control (Hickok, 2012; Hickok et al., 2011), in the current study both tasks also activated frontal areas that are associated with semantic control and domain-general executive processes (Fedorenko et al., 2013; Jackson, 2021). Whilst the executive demands of category fluency are clear (as discussed in detail below), executive control may also important for picture naming in order to resolve which legitimate word to use for each concept (e.g., the hyponym problem: (Levelt, 1992)) as well as competition between the target vs. phonologically- and semantically-related words (G. I. de Zubicaray, Wilson, McMahon, & Muthiah, 2001; G. de Zubicaray, McMahon, Eastburn, & Pringle, 2006). Therefore, the engagement of executive control mechanisms could be regarded as a core component of the speech production network.

### The ATL

The current data also highlight the importance of the ATL as a core-component of the speech production system. Reliable ATL activation was absent in our meta-analysis of the existing literature, presumably due to the poor detection of the signal in this region when using conventional (single echo) fMRI. Indeed, as per other examinations of ATL semantic function (Lambon Ralph et al., 2017), when using fMRI acquisition with improved signal detection then strong ATL activation was found across tasks (Figure 2). Accordingly, this finding helps to realign the neuroimaging literature with evidence from patient studies, whereby damage to the ATL is associated with severe anomia, as well as a generalised semantic impairment across receptive and expressive domains and multiple modalities. Together these results are consistent with the hypothesis that the ATL acts a representational hub for transmodal and transtemporal semantic representation (Jackson, Rogers, & Lambon Ralph, 2021; Lambon Ralph et al., 2017). Indeed, convergent evidence to support the semantic hub hypothesis is found across methodological techniques. For instance, when using cortical grid-electrodes, stimulation of the ATL in patients is associated with naming errors (Rogers et al., 2021; Shimotake et al., 2014) and electrocorticography (ECoG) data have shown ATL semantic coding using multivariate methods (Rogers et al., 2021). Additionally, transcranial magnetic stimulation (TMS) of the ATL impairs responses in naming and comprehension (Pobric et al., 2010; Woollams et al., 2017).

Although both fluency and naming tasks engaged the ATL, the ATL were more strongly engaged by picture naming. Perhaps this is not surprising given that the ATL would be engaged by at least two form of semantic retrieval in the picture naming task – first by processes associated with visual object recognition, and then by processes associated with name retrieval. Indeed, during picture naming there are numerous and extended top-down and bottom-up interaction between the ATL and visual areas, and this elongated process presumably leads to boosted ATL activation (Chan et al., 2011). Actually, the fact that we find ATL activation for verbal fluency at all could be considered surprising, given the ATLs’ anatomical location at the most anterior section of the ventral visual stream, and hence the common suggestion that this region is primarily involved in visual recognition (Duchaine & Yovel, 2015; Kravitz et al., 2011; Skipper, Ross, & Olsoh, 2011).

### The AG

Prominent models propose that the AG functions as a semantic hub that stores semantic information (Binder et al., 2009). In the current study we found no evidence to support this suggestion in either the meta-analysis or fMRI study. Indeed, unlike the ATL and the frontal regions, the AG was systematically deactivated by all tasks relative to rest, with greater or equal levels of deactivation for the semantic tasks relative to the control condition. This is consistent with other speech production studies where the AG is not reliably activated by semantic speech production tasks (Humphreys, Halai, et al., 2022), and where the AG shows greater deactivation for speech production relative to performing non-meaningful tongue movements (Geranmayeh et al., 2012). Parietal damage does not result in gross semantic memory retrieval impairment, unlike damage to the ATL (Jefferies and Lambon Ralph, 2006). The current data add to a growing body of evidence that the AG might not actively contribute to all forms of semantic processing. For instance, direct contrasts have shown that whilst the ATL is positively engaged by semantic tasks, the AG is equally deactivated for semantic and non-semantic control tasks (consistent with AGs role as part of the default mode network (DMN) (Buckner, Andrews-Hanna, & Schacter, 2008)). Furthermore, when processing matched semantic and non-semantic tasks, the AG shows a main effect of task difficulty (easy > hard) but no semantic vs. non-semantic difference (Humphreys & Lambon Ralph, 2017; Quillen, Yen, & Wilson, 2021) and compellingly, classic examples of apparent semantic effects (word > non-word, concrete > abstract) can be flipped by reversing the difficulty of the task or the stimuli (Graves, Boukrina, Mattheiss, Alexander, & Baillet, 2017; Pexman, Hargreaves, Edwards, Henry, & Goodyear, 2007). Finally, whilst the AG is deactivated by many cognitive domains including semantic, there are certain tasks that do positively engage this region (e.g., episodic memory retrieval) (Humphreys, Halai, et al., 2022; Humphreys, Jackson, & Lambon Ralph, 2020; Humphreys, Jung, & Lambon Ralph, 2022; Humphreys & Lambon Ralph, 2015). Together these results suggest that the AG is not a core component of the speech production network, and instead serves alternative functions (Buckner & Carroll, 2007; Humphreys & Lambon Ralph, 2015; Humphreys, Lambon Ralph, & Simons, 2021; Humphreys & Tibon, 2022; Wagner, Shannon, Kahn, & Buckner, 2005).

### Task-specific variance in the extent of lateralisation

Despite an overall bilateral speech production network, the extent of lateralisation was found to be task dependent, with fluency showing boosted left hemisphere engagement compared to naming, particularly in frontal areas. A simple comparison of standard (multi-word) fluency vs. picture (single) naming might suggest that the extent of left-hemisphere bias is dependent on the extent to which a task involved rapid and fluid speech production and complex articulatory coordination. Such differences are unlikely to explain the current results since the two tasks were matched for speech rate and the number of words produced. Furthermore, the result is consistent with existing findings where rapid and fluent speech does appear to be the driving force behind task-specific variations in laterality (Blank et al., 2002; Geranmayeh et al., 2012).

Instead, the locus of activation overlapped with regions associated with semantic and domain-general control. Category fluency loads heavily onto executive tasks (Schumacher et al., 2022) and patients with semantic control impairments show larger category fluency deficits compared to naming (Noonan et al., 2010; Robinson et al., 1998). Furthermore, category fluency correlates highly with overall disease severity in neurodegenerative patients (Hodges & Patterson, 1995), and is poor at differentiating between patient groups despite clear variance in their language abilities (Henderson et al., 2022). This is consistent with the notion that category fluency is a multi-faceted task, relying not only on the speech production system but also regions associated with semantic control, working memory, inhibitory control etc. in the verbal and non-verbal domain (Schumacher et al., 2022). Thus one possible reason for the enhanced leftward lateralisation of fluency over naming tasks is that, since the semantic control network is left-lateralised (Jackson, 2021), when a semantic task requires more executive resources the network is increasingly shifted leftwards due to interactions with the semantic control system. Nevertheless, the function of hemispheric specialisation, and moreover what is the driving force behind such functional biases remains an open question and a considerable amount of research is needed to understand this issue.

### Clinical implications

The current findings are also of clinical relevance: 1) In terms of neuropsychological language assessment, whilst category fluency may be considered a quick and easy speech measure, it is far from a pure measure of speech production as failure can occur even without language impairment given into increased executive demands (Noonan et al., 2010; Robinson et al., 1998), indeed, caution should be drawn against the over-reliance on any single assessment of language function given the task dependent nature of performance. 2) In terms of presurgical fMRI assessment, the degree of lateralisation will depend on what production task is selected (Birn et al., 2010; Blank et al., 2002; Geranmayeh et al., 2012; Moore & Price, 1999; Vitali et al., 2005; Zelkowicz et al., 1998). 3) More generally, if using fMRI to explore language functions in patient populations then the task and fMRI acquisition choices are very important for the ability to see the full speech production neural network (Rice et al., 2018; Robson et al., 2014).

## Conclusions

The current study combined the results of a meta-analysis and fMRI study, directly contrasting picture-naming and category fluency tasks. The results suggest that the core speech production network comprises a bilateral fronto-temporal network including executive and motor frontal areas as well as ATL semantic representational regions. The extent of lateralisation in frontal regions for speech production is task-dependent with more executively-demanding speech tasks (in this case category fluency) associated with boosted left hemisphere activation.

## Acknowledgements

This research was supported by an MRC Programme grant to MALR (MR/R023883/1) and an intramural award (MC_UU_00005/18). The authors declare no conflict of interest.

Data availability statement: The data will be made available at http://www.mrc438cbu.cam.ac.uk/publications/opendata

## References

1. Ashburner, J. (2007). A fast diffeomorphic image registration algorithm. Neuroimage, 38(1), 95–113. doi: DOI 10.1016/j.neuroimage.2007.07.007

2. Binder, J. R., Desai, R. H., Graves, W.W., & Conant, L. (2009). Where Is the Semantic System? A Critical Review and Meta-Analysis of 120 Functional Neuroimaging Studies. Cerebral Cortex, 19(12), 2767–2796. doi: 10.1093/cercor/bhp055

3. Binney, Richard J., Embleton, Karl V., Jefferies, Elizabeth, Parker, Geoffrey J. M., & Lambon Ralph, Matthew A. (2010). The Ventral and Inferolateral Aspects of the Anterior Temporal Lobe Are Crucial in Semantic Memory: Evidence from a Novel Direct Comparison of Distortion-Corrected fMRI, rTMS, and Semantic Dementia. Cerebral Cortex, 20(11), 2728–2738. doi: 10.1093/cercor/bhq019

4. Birn, R. M., Kenworthy, L., Case, L., Caravella, R., Jones, T. B., Bandettini, P.A., & Martin, A. (2010). Neural systems supporting lexical search guided by letter and semantic category cues: a self-paced overt response fMRI study of verbal fluency. Neuroimage, 49(1), 1099–1107.

5. Blank, S. C., Scott, S. K., Murphy, K., Warburton, E., & Wise, R. J. S. (2002). Speech production: Wernicke, Broca and beyond. Brain, 125, 1829–1838.

6. Buckner, R. L., Andrews-Hanna, J. R., & Schacter, D. L. (2008). The brain’s default network - Anatomy, function, and relevance to disease. Year in Cognitive Neuroscience 2008, 1124, 1–38. doi: DOI 10.1196/annals.1440.011

7. Buckner, R. L., & Carroll, D. C. (2007). Self-projection and the brain. Trends in Cognitive Sciences, 11(2), 49–57. doi: 10.1016/j.tics.2006.11.004

8. Chan, Alexander M., Baker, Janet M., Eskandar, Emad, Schomer, Donald, Ulbert, Istvan, Marinkovic, Ksenija, . . . Halgren, Eric. (2011). First-Pass Selectivity for Semantic Categories in Human Anteroventral Temporal Lobe. The Journal of Neuroscience, 31(49), 18119–18129. doi: 10.1523/jneurosci.3122-11.2011

9. Chang, Y., & Lambon Ralph, M. A. (2020). A unified neurocomputational bilateral model of spoken language production in healthy participants and recovery in poststroke aphasia. Proceedings of the National Academy of Sciences, 117(51), 32779–32790. doi: doi:10.1073/pnas.2010193117

10. Chiou, R., Jefferies, E., Duncan, J., Humphreys, G. F., & Lambon Ralph, M. A. (2023). A middle ground where executive control meets semantics: The neural substrates of semantic-control are topographically sandwiched between the multiple-demand and default-mode systems. Cerebral Cortex, 33(8), 4512– 4526. doi: 10.1101/2021.11.26.470178

11. de Zubicaray, Greig I., Wilson, Stephen J., McMahon, Katie L., & Muthiah, Santhi. (2001). The semantic interference effect in the picture-word paradigm: An event-related fMRI study employing overt responses. Human Brain Mapping, 14(4), 218–227. doi: 10.1002/hbm.1054

12. de Zubicaray, Greig, McMahon, Katie, Eastburn, Mathew, & Pringle, Alan. (2006). Top-down influences on lexical selection during spoken word production: A 4T fMRI investigation of refractory effects in picture naming. Human Brain Mapping, 27(11), 864–873. doi: 10.1002/hbm.20227

13. DiCarlo, J. J., & Cox, D. D. Untangling invariant object recognition. Trends in Cognitive Sciences, 11(8), 333–341.

14. Duchaine, B., & Yovel, G. . (2015). A revised neural framework for face processing. Annual Review of Vision Science, 1, 393–416.

15. Eggert, H. (1977). Wernicke’s Works on Aphasia. A Sourcebook and Review. . The Hague, The Netherlands: Mouton.

16. Eickhoff, S. B., Laird, A. R., Grefkes, C., Wang, L. E., Zilles, K., & Fox, P. T. (2009). Coordinate-based activation likelihood estimation meta-analysis of neuroimaging data: a random-effects approach based on empirical estimates of spatial uncertainty. Human Brain Mapping, 30(9), 2907–2926. doi: 10.1002/hbm.20718

17. Fedorenko, E., Duncan, J., & Kanwisher, N. (2013). Broad domain generality in focal regions of frontal and parietal cortex. Proc Natl Acad Sci U S A, 110(41), 16616–16621. doi: 1315235110 [pii] 10.1073/pnas.1315235110

18. Friederici, Angela D. (2002). Towards a neural basis of auditory sentence processing. Trends in Cognitive Sciences, 6(2), 78–84.

19. Gajardo-Vidal, Andrea, Lorca-Puls, Diego L, Hope, Thomas M H, Parker Jones, Oiwi, Seghier, Mohamed L, Prejawa, Susan, . . . Price, Cathy J. (2018). How right hemisphere damage after stroke can impair speech comprehension. Brain, 141(12), 3389–3404. doi: 10.1093/brain/awy270

20. Geranmayeh, F., Brownsett, S. L. E., Leech, R., Beckmann, C. F., Woodhead, Z., & Wise, R. J. S. (2012). The contribution of the inferior parietal cortex to spoken language production. Brain and Language, 121(1), 47–57.

21. Geschwind, N. (1965). Disconnexion syndromes in animals and man. Brain, 88(2), 237–294.

22. Geschwind, N. (1972). Language and Brain. Scientific American, 226(4), 76-&.

23. Graves, W. W., Boukrina, O., Mattheiss, S. R., Alexander, E. J., & Baillet, S. (2017). Reversing the Standard Neural Signature of the Word-Nonword Distinction. Journal of Cognitive Neuroscience, 29(1), 79–94.

24. Hagoort, Peter. (2005). On Broca, brain, and binding: a new framework. Trends in Cognitive Sciences, 9(9), 416–423.

25. Halai, A. D., Welbourne, S. R., Embleton, K., & Parkes, L. M. (2014). A comparison of dual gradient-echo and spin-echo fMRI of the inferior temporal lobe. Human Brain Mapping, 35(8), 4118–4128. doi: 10.1002/hbm.22463

26. Hartwigsen, Gesa, Saur, Dorothee, Price, Cathy J., Ulmer, Stephan, Baumgaertner, Annette, & Siebner, Hartwig R. (2013). Perturbation of the left inferior frontal gyrus triggers adaptive plasticity in the right homologous area during speech production. Proceedings of the National Academy of Sciences, 110(41), 16402–16407. doi: doi:10.1073/pnas.1310190110

27. Henderson, Shalom K, Peterson, Katie A, Patterson, Karalyn, Lambon Ralph, M. A., & Rowe, James. (2022). Verbal fluency tests assess global cognitive status but have limited diagnostic differentiation: Evidence from a large-scale examination of six neurodegenerative diseases. medRxiv, 2022.2008.2016.22278837. doi: 10.1101/2022.08.16.22278837

28. Hickok, G. (2012). Computational neuroanatomy of speech production. Nature Reviews Neuroscience, 13(2), 135–145.

29. Hickok, G., Houde, J., & Rong, F. (2011). Sensorimotor Integration in Speech Processing: Computational Basis and Neural Organization. Neuron, 69(3), 407–422.

30. Hickok, Gregory, & Poeppel, David. (2000). Towards a functional neuroanatomy of speech perception. Trends in Cognitive Sciences, 4(4), 131–138. doi: 10.1016/s1364-6613(00)01463-7

31. Hickok, Gregory, & Poeppel, David. (2004). Dorsal and ventral streams: a framework for understanding aspects of the functional anatomy of language. Cognition, 92(1-2), 67–99.

32. Hickok, Gregory, & Poeppel, David. (2007). The cortical organization of speech processing. Nat Rev Neurosci, 8(5), 393–402.

33. Hodges, J. R., & Patterson, K. (1995). Is semantic memory consistently impaired early in the course of Alzheimer’s disease? Neuroanatomical and diagnostic implications. Neuropsychologia, 33, 441–459.

34. Hoffman, P. (2019). Reductions in prefrontal activation predict off-topic utterances during speech production. Nature Communications, 10(1), 515.

35. Hoffman, P., Loginova, E., & Russell, A. (2018). Poor coherence in older people’s speech is explained by impaired semantic and executive processes. ELife, 7, 1–29.

36. Humphreys, G. F., Halai, A. D., Branzi, F. M., & Lambon Ralph, M. A. (2022). The angular gyrus is engaged by autobiographical recall not object-semantics, or event-semantics: Evidence from contrastive propositional speech production. *bioRxiv*. doi: 10.1101/2022.04.04.487000

37. Humphreys, G. F., Hoffman, P., Visser, M., Binney, R. J., & Ralph, M. A. L. (2015). Establishing task- and modality-dependent dissociations between the semantic and default mode networks. Proceedings of the National Academy of Sciences of the United States of America, 112(25), 7857–7862. doi: 10.1073/pnas.1422760112

38. Humphreys, G. F., Jackson, R. L., & Lambon Ralph, M. A. (2020). Overarching Principles and Dimensions of the Functional Organization in the Inferior Parietal Cortex. Cerebral Cortex, 30(11), 5639–5653.

39. Humphreys, G. F., Jung, J., & Lambon Ralph, M. A. (2022). The convergence and divergence of episodic and semantic functions across lateral parietal cortex. Cerebral Cortex, 1–18. doi: 10.1093/braincomms/fcaa125

40. Humphreys, G. F., & Lambon Ralph, M. A. (2015). Fusion and fission of cognitive functions in the human parietal cortex. Cerebral Cortex, 25, 3547–3560.

41. Humphreys, G. F., & Lambon Ralph, M. A. (2017). Mapping Domain-Selective and Counterpointed Domain-General Higher Cognitive Functions in the Lateral Parietal Cortex: Evidence from fMRI Comparisons of Difficulty-Varying Semantic Versus Visuo-Spatial Tasks, and Functional Connectivity Analyses. Cerebral Cortex, 27(8), 4199–4212.

42. Humphreys, G. F., Lambon Ralph, M. A., & Simons, J. S. (2021). A unifying account of angular gyrus contributions to episodic and semantic cognition. Trends in Neurosciences, 44(6), 452–463. doi: 10.31234/osf.io/r2deu

43. Humphreys, G. F., & Tibon, R. (2022). Dual-axes of functional organisation across lateral parietal cortex: the angular gyrus forms part of a multi-modal buffering system. Brain Structure & Function. doi: 10.1007/s00429-022-02510-0

44. Indefrey, P. (2011). The spatial and temporal signatures of word production components: a critical update. Frontiers in Psychology, 2, 255.

45. Indefrey, P., & Levelt, W. J. M. (2004). The spatial and temporal signatures of word production components. Cognition, 92, 101–144.

46. Jackson, R. L. (2021). The neural correlates of semantic control revisited. Neuroimage, 224, 117444. doi: 10.1016/j.neuroimage.2020.117444

47. Jackson, R. L., Rogers, T. T., & Lambon Ralph, M. A. (2021). Reverse-engineering the cortical architecture for controlled semantic cognition. Nature Human Behaviour, 5, 774–786.

48. Jefferies, E., & Lambon Ralph, M. A. (2006). Semantic impairment in stroke aphasia versus semantic dementia: a case-series comparison. Brain, 129, 2132–2147. doi: Doi 10.1093/Brain/Awl153

49. Johnstone, T., Ores Walsh, K. S., Greischar, L. L., Alexander, A. L., Fox, A. S., & Davidson, R. J. (2006). Motion correction and the use of motion covariates in multiple-subject fMRI analysis. Human Brain Mapping, 27, 779–788. doi: 10.1002/hbm.20219

50. Jung, J., & Lambon Ralph, M. A. (2016). Mapping the dynamic network interactions underpinning cognition: A cTBS-fMRI study of the flexible adaptive neural system for semantics. Cerebral Cortex, 26, 3580–3590

51. Kravitz, D. J., Saleem, K. S., Baker, C. I., & Mishkin, M. (2011). A new neural framework for visuospatial processing. Nature Reviews Neuroscience, 12(4), 217–230. doi: Doi 10.1038/Nrn3008

52. Krieger-Redwood, K., & Jefferies, E. (2014). TMS interferes with lexical-semantic retrieval in left inferior frontal gyrus and posterior middle temporal gyrus: Evidence from cyclical picture naming. Neuropsychologia, 64, 24–32.

53. Laird, A. R., Fox, P. M., Price, C. J., Glahn, D. C., Uecker, A. M., Lancaster, J. L., . . . Fox, P. T. (2005). ALE meta-analysis: controlling the false discovery rate and performing statistical contrasts. Human Brain Mapping, 25(1), 155–164. doi: 10.1002/hbm.20136

54. Lambon Ralph, M. A., Cipolotti, Lisa, Manes, Facundo, & Patterson, Karalyn. (2010). Taking both sides: do unilateral anterior temporal lobe lesions disrupt semantic memory? Brain, 133(11), 3243–3255. doi: 10.1093/brain/awq264

55. Lambon Ralph, M. A., Ehsan, S., Baker, G. A., & Rogers, T. T. (2012). Semantic memory is impaired in patients with unilateral anterior temporal lobe resection for temporal lobe epilepsy. . Brain, 135, 242–258.

56. Lambon Ralph, M. A., Jefferies, E., Patterson, K., & Rogers, T. T. (2017). The neural and computational bases of semantic cognition. Nat Rev Neurosci, 18(1), 42–55. doi: 10.1038/nrn.2016.150

57. Lambon Ralph, M. A., McClelland, J. L., Patterson, K., Galton, C. J., & Hodges, J. R. (2001). No right to speak? The relationship between object naming and semantic impairment: Neuropsychological evidence and a computational model. Journal of Cognitive Neuroscience, 13, 341–356.

58. Lambon Ralph, M. A., Pobric, G., & Jefferies, E. (2009). M. A. Lambon Ralph, G. Pobric, E. Jefferies, Conceptual knowledge is underpinned by the temporal pole bilaterally: Convergent evidence from rTMS. Cerebral Cortex, 19, 832– 838.

59. Lambon Ralph, M. A., Sage, K., Jones, R. W., & Mayberry, E. J. (2010). Coherent concepts are computed in the anterior temporal lobes. Proceedings of the National Academy of Sciences of the United States of America, 107(6), 2717–2722. doi: DOI 10.1073/pnas.0907307107

60. Levelt, W. J. M. (1992). Accessing words in speech production: Stages, processes and representations. Cognition, 42(1), 1–22.

61. Lichtheim, L. (1885). On aphasia. Brain, 7, 433–484.

62. Moore, C. J., & Price, C. (1999). Three Distinct Ventral Occipitotemporal Regions for Reading and Object Naming. Neuroimage, 10, 181–192.

63. Noonan, K. A., Jefferies, E., Corbett, F., & Lambon Ralph, M. A. (2010). Elucidating the Nature of Deregulated Semantic Cognition in Semantic Aphasia: Evidence for the Roles of Prefrontal and Temporo-parietal Cortices. Journal of Cognitive Neuroscience, 22, 1597–1613.

64. Noonan, K. A., Jefferies, E., Visser, M., & Lambon Ralph, M. A. (2013). Going beyond inferior prefrontal involvement in semantic control: evidence for the additional contribution of dorsal angular gyrus and posterior middle temporal cortex. Journal of Cognitive Neuroscience, 25(11), 1824–1850. doi: 10.1162/jocn_a_00442

65. Pexman, P. M., Hargreaves, I. S., Edwards, J. D., Henry, L. C., & Goodyear, B. G. (2007). Neural correlates of concreteness in semantic categorization. Journal of Cognitive Neuroscience, 19(8), 1407–1419. doi: DOI 10.1162/jocn.2007.19.8.1407

66. Pobric, G., Jefferies, E., & Lambon Ralph, M. A. (2010). Amodal semantic representations depend on both anterior temporal lobes: evidence from repetitive transcranial magnetic stimulation. Neuropsychologia, 48, 1336–1342.

67. Quillen, Ian A., Yen, Melodie, & Wilson, Stephen M. (2021). Distinct neural correlates of linguistic demand and non-linguistic demand. Neurobiology of Language, 0(ja), 1–59. doi: 10.1162/nol_a_00031

68. Rice, G. E., Caswell, H., Moore, P., Hoffman, P., & Lambon Ralph, M. A. (2018). The Roles of Left Versus Right Anterior Temporal Lobes in Semantic Memory: A Neuropsychological Comparison of Postsurgical Temporal Lobe Epilepsy Patients. Cerebral Cortex, 28(4), 1487–1501. doi: 10.1093/cercor/bhx362

69. Rice, G. E., Lambon Ralph, M. A., & Hoffman, P. (2015). The roles of left versus right anterior temporal lobes in conceptual knowledge: an ALE meta-analysis of 97 functional neuroimaging studies. Cerebral Cortex, 25(11), 4374–4391.

70. Riesenhuber, M., & Poggio, T. (1999). Hierarchical models of object recognition in cortex. Nature Neuroscience, 2(11), 1019–1025.

71. Robinson, G., Blair, J., & Cipolotti, L. (1998). Dynamic aphasia: an inability to select between competing verbal responses. Brain, 121, 77–89.

72. Robson, Holly, Zahn, Roland, Keidel, James L., Binney, Richard J., Sage, Karen, & Lambon Ralph, Matthew A. (2014). The anterior temporal lobes support residual comprehension in Wernicke’s aphasia. Brain, 137(3), 931–943. doi: 10.1093/brain/awt373

73. Rogers, T. T., Cox, C. R., Lu, Q., Shimotake, A., Kikuchi, T., Kunieda, T., . . . Lambon Ralph, M. A. (2021). Evidence for a deep, distributed and dynamic code for animacy in human ventral anterior temporal cortex. ELife, 10, e66276. doi: 10.7554/eLife.66276

74. Rogers, T. T., Patterson, K., Jefferies, E., & Lambon Ralph, M. A. (2015). Rogers, T. T., Patterson, K., Jefferies, E. & Lambon Ralph, M. A. Disorders of representation and control in semantic cognition: Effects of familiarity, typicality, and specificity. Neuropsychologia, 76, 220–239.

75. Sato, N., Matsumoto, R., Shimotake, A., Matsuhashi, M., Otani, M., Kikuchi, T., . . . Ikeda, A. (2021). Frequency-Dependent Cortical Interactions during Semantic Processing: An Electrocorticogram Cross-spectrum Analysis Using a Semantic Space Model. Cerebral Cortex, 31(9), 4329–4339.

76. Schapiro, A. C., McClelland, J. L., Welbourne, S. R., Rogers, T. T., & Lambon Ralph, M. A. (2013). Why Bilateral Damage Is Worse than Unilateral Damage to the Brain. Journal of Cognitive Neuroscience, 25(12), 2107–2123. doi: 10.1162/jocn_a_00441

77. Schumacher, R., Halai, A. D., & Lambon Ralph, M. A. (2022). Assessing executive functions in post-stroke aphasia—utility of verbally based tests. Brain Communications, 4, fcac107.

78. Shimotake, Akihiro, Matsumoto, Riki, Ueno, Taiji, Kunieda, Takeharu, Saito, Satoru, Hoffman, Paul, . . . Lambon Ralph, Matthew A. (2014). Direct Exploration of the Role of the Ventral Anterior Temporal Lobe in Semantic Memory: Cortical Stimulation and Local Field Potential Evidence From Subdural Grid Electrodes. Cerebral Cortex, 25(10), 3802–3817. doi: 10.1093/cercor/bhu262

79. Skipper, L. M., Ross, L. A., & Olsoh, I. R. (2011). Sensory and semantic category subdivisions within the anterior temporal lobes. Neuropsychologia, 49(12), 3419–3429.

80. Snodgrass, J. G., & Vanderwart, M. (1980). A standardized set of 260 pictures: Norms for name agreement, image agreement, familiarity, and visual complexity. Journal of Experimental Psychology: Human Learning and Memory, 6(2), 174–215.

81. Stefaniak, J. D., Geranmayeh, F., & Lambon Ralph, M. A. (2022). The multidimensional nature of aphasia recovery post-stroke. Brain, 145(4), 1354–1367. doi: 10.1093/brain/awab377

82. Stefaniak, J. D., Halai, A. D., & Lambon Ralph, M. A. (2020). The neural and neurocomputational bases of recovery from post-stroke aphasia. Nature Reviews Neuroscience, 16, 43–55.

83. Ueno, T., Saito, S., Rogers, T. T., & Lambon Ralph, M. A. (2011). Lichtheim 2: Synthesizing Aphasia and the Neural Basis of Language in a Neurocomputational Model of the Dual Dorsal-Ventral Language Pathways. Neuron, 72(2), 385–396. doi: DOI 10.1016/j.neuron.2011.09.013

84. Visser, M., Jefferies, E., & Lambon Ralph, M. A. (2010). Semantic Processing in the Anterior Temporal Lobes: A Meta-analysis of the Functional Neuroimaging Literature. Journal of Cognitive Neuroscience, 22(6), 1083–1094.

85. Vitali, P., Abutalebi, J., Tettamanti, M., Rowe, J., Scifo, P., Fazio, F., . . . Perani, D. (2005). Generating animal and tool names: An fMRI study of effective connectivity. Brain and Language, 93(1), 32–45.

86. Wagner, A. D., Shannon, B. J., Kahn, I., & Buckner, R. L. (2005). Parietal lobe contributions to episodic memory retrieval. Trends in Cognitive Sciences, 9(9), 445–453. doi: DOI 10.1016/j.tics.2005.07.001

87. Weiller, C., Bormann, T., Saur, D., Musso, M., & Rijntjes, M. (2011). How the ventral pathway got lost – And what its recovery might mean. Brain and Language, 118(1), 29–39.

88. Whitney, Carin, Kirk, Marie, O’Sullivan, Jamie, Lambon Ralph, Matthew A., & Jefferies, Elizabeth. (2010). The Neural Organization of Semantic Control: TMS Evidence for a Distributed Network in Left Inferior Frontal and Posterior Middle Temporal Gyrus. Cerebral Cortex, 21(5), 1066–1075. doi: 10.1093/cercor/bhq180

89. Woollams, A., Lindley, L. J., Pobric, G., & Hoffman, P. (2017). Laterality of anterior temporal lobe repetitive transcranial magnetic stimulation determines the degree of disruption in picture naming. Brain Structure & Function, 222, 3749–3759.

90. Zelkowicz, B., Herbster, A., Nebes, R., Mintun, M., & Becker, A. . (1998). An examination of regional cerebral blood flow during object naming tasks. Journal of the International Neuropsychological Society, 4(2), 160–166.

