## Supplementary for "Mapping the task-general and task-specific neural correlates of speech production: meta-analysis and fMRI direct comparisons of category fluency and picture naming"

Table S1. The clusters identified from the meta-analysis of Category Fluency and Picture Naming studies

|  | Cluster # | x | y | z | Z | Anatomical label |
| --- | --- | --- | --- | --- | --- | --- |
| Fluency studies | 1 | -44 | 24 | 24 | 8.489372 | Left Middle frontal gyrus |
|  | 1 | -42 | 8 | 26 | 6.206844 | Left Inferior frontal gyrus |
|  | 1 | -46 | -12 | 36 | 5.598102 | Left Precentral gyrus |
|  | 2 | -6 | 12 | 44 | 10.21135 | Left Supplementary Motor Area |
|  | 3 | -24 | -2 | 50 | 6.453395 | Left Middle frontal gyrus |
|  | 4 | -14 | -8 | 16 | 5.2552 | Left Thalamus |
|  | 5 | -28 | -58 | 32 | 3.411601 | Left Middle Temporal Gyrus |
|  | 6 | 48 | -10 | 38 | 4.959919 | Right Precentral gyrus |
|  | 7 | -36 | -36 | -2 | 4.334408 | Left Caudate |
|  | 8 | 42 | 10 | 32 | 3.678642 | Right Precentral gyrus |
|  | 9 | 40 | 26 | -10 | 3.606574 | Right Inferior frontal gryus |
|  | 10 | 36 | -62 | -28 | 3.189645 | Right Cerebellum |
|  | 11 | 40 | 36 | 16 | 3.146989 | Right Middle Frontal Gyrus |
|  | 12 | -2 | 2 | 68 | 3.204904 | Left Superior Frontal Gyrus |
| Naming studies | 1 | -44 | -60 | -16 | 7.750345 | Left Fusiform Gyrus |
|  | 1 | -36 | -44 | -16 | 7.008279 | Left Fusiform Gyrus |
|  | 1 | -38 | -82 | -8 | 4.462598 | Left Fusiform Gyrus |
|  | 2 | -48 | 10 | 8 | 6.025524 | Left Precentral gyrus |
|  | 2 | -46 | 28 | 2 | 5.835731 | Left Inferior frontal gyrus |
|  | 2 | -44 | 8 | 28 | 5.772944 | Left Inferior frontal gyrus |
|  | 2 | -48 | 18 | 24 | 5.729263 | Left Middle frontal gyrus |
|  | 2 | -34 | 24 | 2 | 4.468136 | Left Insula |
|  | 3 | 32 | -48 | -14 | 4.616426 | Right Cerebellum |
|  | 4 | -6 | 8 | 54 | 4.353867 | Left Supplementary Motor Area |
|  | 5 | 34 | -88 | 10 | 4.809121 | Right Occipital lobe |
|  | 6 | -52 | -4 | 48 | 5.190566 | Left Precentral gyrus |
|  | 7 | -28 | -96 | 8 | 4.692374 | Left Occipital Lobe |
|  | 8 | -54 | -30 | 0 | 3.677164 | Left Middle Temporal Gyrus |
|  | 9 | 58 | -4 | 38 | 3.611051 | Right Precentral gyrus |

Table S2. The peak coordinates of activation from the fMRI study.

| Uncorrected p < .001, cluster corrected | | | | | |
| --- | --- | --- | --- | --- | --- |
| Contrast | t | x | y | z | Anatomical label |
| Fluency > control | 18.03 | -3 | 14 | 48 | cingulate gyrus/motor cortex |
|  | 15.36 | -5 | 20 | 38 | paracingulate gyrus |
|  | 15.18 | 13 | -77 | -29 | Cerebellum |
|  | 14.14 | -30 | 27 | -3 | Orbito-frontal cortex |
|  | 13.71 | 20 | 25 | 31 | Cingulate gyrus |
|  | 13.41 | -30 | 22 | 3 | Insular |
|  | 12.86 | -39 | -3 | 51 | Precentral gyrus |
|  | 12.11 | -42 | 21 | 25 | inferior frontal gyrus, BA44 |
|  | 11.67 | 12 | 16 | 37 | Cingulate gyrus |
|  | 10.22 | -26 | -74 | 35 | Superior parietal lobule |
|  | 10.1 | 36 | 18 | 3 | Insular |
|  | 9.41 | 36 | 21 | -13 | Orbito-frontal cortex |
|  | 8.82 | 16 | 4 | 19 | Caudate |
|  | 8.37 | -9 | -72 | 7 | Visual cortex |
|  | 7.86 | -4 | -8 | 6 | Thalamus |
|  | 7.83 | -14 | -3 | 14 | Caudate |
|  | 7.76 | -30 | 60 | 6 | Frontal pole |
|  | 7.13 | 52 | -12 | 41 | Postcentral gyrus |
|  | 6 | -36 | -25 | -26 | Fusiform gyrus |
|  | 5.47 | 38 | -21 | -25 | Fusiform gyrus |
| Naming > control | 17.28 | 36 | -61 | -16 | Fusiform gyrus |
|  | 16.96 | 46 | -71 | 17 | Lateral occipital cortex |
|  | 16.67 | -2 | 8 | 48 | Cingulate gyrus/motor cortex |
|  | 15.4 | 34 | -47 | -21 | Fusiform gyrus |
|  | 14.54 | -35 | -85 | -9 | Lateral occipital cortex |
|  | 14.24 | -33 | -69 | -18 | Fusiform gyrus |
|  | 13.96 | -36 | -73 | -18 | Fusiform gyrus |
|  | 12.16 | -28 | -74 | 26 | Lateral occipital cortex |
|  | 11.56 | -29 | 26 | 1 | Insular |
|  | 10.62 | 54 | -11 | 42 | Postcentral gyrus |
|  | 10.11 | -43 | -13 | 42 | Middle frontal gyrus |
|  | 9.95 | 40 | -21 | -25 | Fusiform gyrus |
|  | 8.3 | -39 | 20 | 25 | inferior frontal gyrus, BA44 |
|  | 8.3 | -31 | -58 | 55 | Superior parietal lobule |
|  | 7.57 | -36 | -27 | -25 | Fusiform gyrus |
|  | 7.25 | -47 | 6 | 38 | inferior frontal gyrus, BA44 |
| Fluency > naming | 11.48 | -6 | 15 | 48 | Cingulate gyrus/premotor cortex |
|  | 10.75 | -27 | 12 | 48 | Middle frontal gyrus |
|  | 9.85 | -3 | 24 | 36 | Cingulate gyrus |
|  | 9.04 | 15 | -87 | -39 | Cerebellum |
|  | 8.97 | 12 | 27 | 30 | Cingulate gyrus |
|  | 8.71 | -24 | 54 | 0 | Frontal pole |
|  | 8.63 | -45 | 27 | 24 | Middle frontal gyrus |
|  | 8.29 | 12 | -81 | -30 | Cerebellum |
|  | 7.74 | 33 | 36 | 30 | Frontal pole |
|  | 7.52 | -3 | 12 | 63 | Superior frontal gyrus |
|  | 7.32 | 27 | 9 | 57 | Middle frontal gyrus |
|  | 7.04 | -9 | 33 | 18 | Anterior cingulate cortex |
|  | 6.86 | 0 | 30 | 15 | Anterior cingulate cortex |
|  | 6.56 | 9 | 36 | 15 | Anterior cingulate cortex |
|  | 6.52 | -36 | -78 | 42 | Angular gyrus, PGp |
|  | 5.92 | -24 | 33 | 30 | Middle frontal gyrus |
|  | 5.342 | 21 | 3 | 69 | Premotor cortex |
|  | 5.34 | -15 | 63 | 15 | Frontal pole |
|  | 5.17 | -3 | 39 | 51 | Superior frontal gyrus |
|  | 4.60 | -48 | 30 | 3 | Inferior frontal gyrus, BA45 |
| Naming > fluency | 22.43 | 36 | -57 | -15 | Fusiform gyrus |
|  | 21.82 | 39 | -69 | -12 | Lateral occipital cortex |
|  | 19.72 | -36 | -84 | -6 | Lateral occipital cortex |
|  | 18.81 | -39 | -78 | -12 | Lateral occipital cortex |
|  | 18.77 | -33 | -90 | 12 | Occipital pole |
|  | 17.93 | 36 | -84 | 6 | Lateral occipital cortex |
|  | 16.47 | 33 | -84 | 15 | Fusiform gyrus |
|  | 15.45 | 30 | -72 | 30 | Lateral occipital cortex |
|  | 10.2944 | 30 | -57 | 57 | Superior parietal lobule |
|  | 10.15 | 30 | -54 | 51 | Superior parietal lobule/Intra-parietal sulcus |
|  | 8.15 | 42 | 6 | 27 | Precentral gyrus |
|  | 7.67 | 51 | 33 | 9 | Inferior frontal gyrus, BA45 |
|  | 6.39 | -36 | -6 | 15 | Insular |
|  | 6.06 | -66 | -27 | 30 | Supramarginal gyrus |
|  | 6.05 | -66 | -15 | 30 | Postcentral gyrus |
|  | 5.95 | 24 | -3 | -12 | Amygdala |
|  | 5.93 | 66 | -21 | 36 | Supramarginal gyrus |
|  | 5.76 | 36 | -9 | 12 | Insular |
|  | 5.69 | 24 | 0 | -6 | Putamen |
|  | 5.57 | 33 | -9 | -42 | Fusiform gyrus |
|  | 5.54 | -24 | -3 | -15 | Amygdala |
|  | 5.49 | -57 | 0 | 33 | Precentral gyrus |
|  | 5.36 | 18 | -30 | -3 | Thalamus |
|  | 5.31 | -30 | -54 | 51 | Superior parietal lobule |
|  | 5.24 | -24 | -63 | 48 | Superior parietal lobule |
|  | 5.14 | -27 | -3 | -3 | Putamen |
|  | 5.00 | 27 | 3 | 3 | Putamen |
|  | 4.61 | -51 | -30 | 57 | Postcentral gyrus |
